# Reversible Mitochondrial Fragmentation in iPSC-Derived Cardiomyocytes from Children with DCMA, a Mitochondrial Cardiomyopathy

**DOI:** 10.1101/732339

**Authors:** Leili Rohani, Pranav Machiraju, Rasha Sabouny, Guoliang Meng, Shiying Liu, Tian Zhao, Fatima Iqbal, Amir Ravandi, Joseph C. Wu, Aneal Khan, Timothy Shutt, Derrick Rancourt, Steven C. Greenway

## Abstract

**Background:** The dilated cardiomyopathy with ataxia syndrome (DCMA) is an understudied autosomal recessive disease caused by loss-of-function mutations in the poorly characterized gene *DNAJC19*. Clinically, DCMA is commonly associated with heart failure and early death in affected children through an unknown mechanism. DCMA has been linked to Barth syndrome, a rare but well-studied disorder caused by deficient maturation of cardiolipin (CL), a key mitochondrial membrane phospholipid.

**Methods:** Peripheral blood mononuclear cells from two children with DCMA and severe cardiac dysfunction were reprogrammed into induced pluripotent stem cells (iPSCs). Patient and control iPSCs were differentiated into beating cardiomyocytes (iPSC-CMs) using a metabolic selection strategy and mitochondrial structure and CL content before and after incubation with the mitochondrially-targeted peptide SS-31 were quantified.

**Results:** Patient iPSCs carry the causative *DNAJC19* mutation (rs137854888) found in the Hutterite population and the iPSC-CMs demonstrated highly fragmented and abnormally-shaped mitochondria associated with an imbalanced isoform ratio of the mitochondrial protein OPA1, an important regulator of mitochondrial fusion. These abnormalities were reversible by incubation with SS-31 for 24 hours. Differentiation of iPSCs into iPSC-CMs increased the number of CL species observed but consistent, significant differences in CL content were not seen between patients and control.

**Conclusions:** We describe a unique and novel cellular model that provides insight into the mitochondrial abnormalities present in DCMA and identifies SS-31 as a potential therapeutic for this devastating disease.

## Introduction

The dilated cardiomyopathy with ataxia syndrome (DCMA) is an autosomal recessive disorder characterized clinically by early-onset cardiac and neurological features (i.e. dilated cardiomyopathy, prolongation of the QT interval, hypotonia, cerebellar ataxia, developmental delay) with 3-methylglutaconic aciduria and other variable systemic features (e.g. abnormal male genitalia, growth failure, microcytic anemia, hepatic steatosis).^1^ DCMA is highly prevalent in the Canadian Dariusleut Hutterite population who represent the largest collection of affected individuals in the world.^2^ The Hutterites are a religious group, originally immigrating to North America from Europe, that are genetically isolated and share a communal environment. In the Hutterites, DCMA is caused by a single intronic pathogenic variant NG_022933.1:c.130-1G>C (rs137854888) in the *DNAJC19* gene resulting in abnormal splicing and a truncated, non-functional protein.^2^ *DNAJC19* is an inner mitochondrial membrane (IMM) protein and, although some of its interacting partners have been identified,^3^ its precise role is unknown and the underlying mechanism by which the *DNAJC19* mutation contributes to the DCMA phenotype remains elusive. End-stage heart failure and death in early childhood are common in DCMA patients and, until recently, no effective therapeutic had been identified.^4^ Due to the rarity of the disease, randomized clinical drug trials are infeasible and no cellular or animal models have been published to date.

Patient-specific iPSCs have attracted interest for their advantages in elucidating the mechanism underlying rare genetic diseases.^5–8^ Although cell culture models have started to elucidate a role for DNAJC19 as a regulator of IMM structure and function, no one has studied this function in the context of a pathogenic DNAJC19 mutation.^3^ Due to the presence of shared biochemical abnormalities and phenotypic overlap, DCMA (3-methylglutaconic aciduria type V) has been grouped together with Barth syndrome (3-methylglutaconic aciduria type II), a rare X-linked disorder that arises from mutations in *tafazzin* (TAZ) leading to deficiencies in cardiolipin (CL) maturation causing cardiomyopathy (left ventricular dilation or non-compaction), skeletal myopathy, growth retardation and neutropenia.^9^ As a key component of mitochondrial membranes, CL has an important role in the proper formation of mitochondrial protein complexes and is disrupted in heart failure.^10^ The potential for abnormal mitochondrial structure and function in DCMA due to impaired CL remodeling^3^ represents a possible target for therapeutic intervention with the mitochondrially-targeted peptide SS-31 (also known as elamipretide or Bendavia) that interacts specifically with CL to affect membrane curvature, prevent peroxidative damage and maintain mitochondrial structure and integrity.^11–13^

Here, we characterize DCMA-specific iPSCs and the resulting cardiomyocytes generated from two pediatric Hutterite patients with severe cardiac dysfunction. Using these unique and novel cells, we quantified CL composition and mitochondrial structure and evaluated the effect of SS-31.

## Material and Methods

### Generation and Characterization of DCMA iPSCs

This study was approved by the Conjoint Health Research Ethics Board at the University of Calgary and the Research Ethics Board at Stanford University. Written, informed consent was obtained from each participating family. Control and DCMA iPSCs were reprogrammed from peripheral blood mononuclear cells (PBMCs) using Sendai virus according to the protocol developed in the laboratory of Dr. Joseph Wu and the Stanford Cardiovascular Institute Biobank.^14^ The generated iPSCs were fully characterized using standard pluripotency tests.^15^

### Reverse Transcription Polymerase Chain Reaction (RT-PCR) and Genomic DNA (gDNA) Sequencing of iPSCs

To confirm the presence of the *DNAJC19* mutation, iPSCs were collected for RNA isolation once they reached 70-80% confluency. Total RNA was extracted using the PureLink RNA Mini Kit (ThermoFisher Scientific) according to the manufacturer’s protocol followed by DNAse I (ThermoFisher Scientific) digestion. Synthesis of cDNA was performed using RNA (1 µg), Oligo(dT)20 primers (50 µM) and the Superscript IV Reverse Transcriptase kit (ThermoFisher Scientific). PCR amplification was performed in a final volume of 50 μL using Taq DNA Polymerase (ThermoFisher Scientific) and consisted of the following steps: 96°C for 2 min, followed by 35 cycles of 96°C for 30 sec, 56°C for 30 sec, 72°C for 1 min, and 72°C for 7 min. The DNAJC19-specific primers RTF and RTR were designed to amplify a 525 bp cDNA product corresponding to the full-length coding sequence including the 5’ and 3’ UTR boundaries. The housekeeping gene β-actin was used for normalization. For gDNA sequencing, DNA was extracted from iPSCs using phenol:chloroform (OmniPur). PCR amplification of the *DNAJC19* gene was performed in four amplicons: segment 1 (exon 6), segment 2 (exons 4 & 5), segment 3 (exons 1, 2, and 3) and segment 4 (intron 3). All primer sequences are listed in Supplemental Table S1. Amplified products were separated on a 1.5% agarose gel, stained with ethidium bromide and then visualized and photographed on a UV transilluminator. All cDNA and gDNA PCR products were sequenced in the DNA Services Laboratory at The University of Calgary.

### Differentiation of iPSCs into Cardiomyocytes

Both control and DCMA iPSCs were differentiated into cardiomyocytes based on modifications to a published protocol.^16^ Briefly, iPSC colonies were cultured on plates coated with Matrigel (BD Biosciences, cat. BD354277) and cultured in mTeSR1 medium (STEMCELL Technologies, cat. 85875) until they were 80-90% confluent. Differentiation was initiated by changing to RPMI1640 medium containing B-27 (minus insulin) and the glycogen synthase kinase-3 (GSK3) inhibitor CHIR99021 (STEMCELL Technologies, cat. 72052) for 24 h. Cells were exposed to the Wnt signaling inhibitor IWP-4 (STEMCELL Technologies, cat. 72552) for 2 days from days (D) 3 to 5 of differentiation. The cells were then cultured in RPMI1640 medium containing B-27 (minus insulin) for 3 days (D5-D8). A cardiomyocyte purification protocol using L-lactic acid was applied from D8-D12 in glucose-depleted media.^17^ Cardiomyocyte clusters were transferred onto gelatin-coated plates containing DMEM/F12 medium supplemented with KnockOut serum replacement (ThermoFisher Scientific, cat. 10828010), GlutaMAX (ThermoFisher Scientific, cat. 35050061) and non-essential amino acids (ThermoFisher Scientific, cat. 11140050) on D12 and maintained in the same media until D20 of differentiation. Media was changed every 3-4 days. Mature iPSC-CMs were collected at either day 20 or 21 of differentiation and were used for all analyses.

### Flow Cytometry

To assess the efficiency of our cardiac differentiation protocol, flow cytometry was performed using the cardiac-specific marker troponin T2 (TNNT2). The iPSC-CMs were treated with 0.25% Trypsin-EDTA solution (ThermoFisher Scientific, cat. 25200114) for 5-10 min and then dissociated into single cells. Cells were washed with cardiomyocyte maintenance medium and then centrifuged at 200*g* for 5 min. Cells were then resuspended in 2% paraformaldehyde (PFA, J.T. Baker, cat. 2106-01) and fixed at room temperature (RT) for 15 min. Fixed cells were then washed three times in 4 mL of Dulbecco’s phosphate-buffered saline (DPBS) and permeabilized using 0.2% Triton X-100 (Sigma-Aldrich, cat. T8787) in DPBS at 37°C for 20 min. Finally, cells were washed with flow buffer containing 0.5% bovine serum albumin (BSA) in PBS (ThermoFisher Scientific, cat. 37525) and then incubated in 10% BSA solution at 37°C for 30 min. The primary antibody TNNT2 (ThermoFisher Scientific, cat. MS295-P1) was diluted 1:2000 in flow buffer and incubated with the cells for 60 min at RT. After washing twice with flow buffer, cells were incubated with secondary antibody AlexaFluor 488 (ThermoFisher Scientific, cat. A21121), diluted 1:2000 in flow buffer, for 30 min at RT. Cells were then washed and resuspended in 200 µL flow buffer within a FACS tube and analyzed using a BD FACSVantage SE System in the University of Calgary Flow Cytometry Core Facility. Doublet discrimination gating was performed and gates were drawn based on isotype controls. The histogram of flow cytometry analysis was generated by FlowJo V10.1.3 software.

### Immunocytochemistry

Following differentiation, immunocytochemistry for TNNT2 and NKX2.5 was performed using the Cardiomyocyte Immunocytochemistry kit (ThermoFisher Scientific, cat. A25973). Nuclei were counterstained using NucBlue and stained cells were imaged using a Zeiss Axiovert microscope with a 20x objective. To visualize the mitochondrial network, iPSC-CMs were stained with TOMM20. Prior to staining, cells were plated as single cells (5 × 10^5^ cells) on 35 mm FluoroDish Cell Culture Dishes (World Precision Instruments, cat. WD35-100) and allowed to attach for 72 h. For the SS-31 treated group, cells were incubated with 100 nM SS-31 (d-Arg-2’,6’-dimethyltyrosine-Lys-Phe-NH_2_, synthesized for us by China Peptides) or vehicle control for 24 h. Cells were washed twice with DPBS and then fixed with 4% PFA in DPBS at 37°C for 15 min. Cells were then washed three times with DPBS and quenched with 50 mM NH_4_Cl for 15 min at RT followed by washing with DPBS. Cells were permeabilized with 0.2% Triton X-100 in DPBS for 15 min followed by blocking with 10% FBS for 25 min at RT. The cells were incubated with TOMM20 primary antibody (Sigma-Aldrich, cat. HPA011562), diluted 1:1000 in 5% fetal bovine serum (FBS), for 1 h at 37°C and then washed three times. Secondary antibody AlexaFluor 488 was diluted 1:1000 in 5% FBS and added to the cells for 1 h at RT. Cells were then washed and stored at 4ºC in the dark until imaged on a Zeiss LSM880 confocal microscope using a 63x oil objective.

### Quantification of Mitochondrial Network Fragmentation

Thirty cells from each control and DCMA group were randomly selected and manually graded by a blinded researcher using a published scoring system.^18^

### Transmission Electron Microscopy (TEM)

To assess mitochondrial ultrastructure, control and DCMA iPSC-CMs were cultured on gelatinized plastic 35 mm cell-culture treated plates (5 × 10^5^ cells) for three days. For the SS-31 treated group, cells were incubated with 100 nM SS-31 or vehicle control for 24 h. Cells were then fixed and imaged using a Hitachi H7650 microscope in The University of Calgary’s Microscopy and Imaging Facility. TEM images were exported into FIJI (ImageJ) and 10 mitochondrion were analyzed per group using the “measure perimeter” function.

### Western Blotting

We measured the long (L) and short (S) forms of OPA1 using western blotting. All iPSC-CMs were cultured on gelatinized plastic 35 mm cell-culture treated plates (5 × 10^5^ cells) for 72 h. For the SS-31 treated group, cells were incubated with 100 nM SS-31 or vehicle control for 24 h. Subsequently, cells were harvested, washed and lysed with RIPA buffer containing protease inhibitors (Amersco, cat. M250). Total cell lysates (20 μg) were resolved by SDS-PAGE and transferred onto PVDF membrane. Blots were probed with antibodies against OPA1 (BD Bioscience, cat. 612606) and loading control HSP60 (Cell Signaling, cat. 12165) at a 1:1000 final dilution followed by horseradish peroxidase-conjugated secondary antibodies. Blots were incubated with Clarity ECL substrate (BioRad) according to the manufacturer’s instructions and imaged on an Amersham Imager AI600. Quantitation of protein band intensities was performed using ImageJ and normalized to HSP60. Data are represented as the ratio of L-OPA1 / S-OPA1 from three independent experiments.

### Cardiolipin Quantification

Mass spectrometry was used to quantify the CL of control and DCMA iPSCs and iPSC-CMs as previously described.^19^ Two separate preparations from iPSCs and three separate preparations from iPSC-CMs cultured for 21 days were analyzed.

### Statistical Analysis

Data analysis was performed in Prism 8 (GraphPad Software). All data are expressed as mean ± standard deviation (SD) or standard error of the mean (SEM). Significance was determined using a two-way ANOVA with Holm-Sidak multiple-testing correction for the mitochondrial fragmentation data after normalization. A one-way ANOVA was used with Holm-Sidak multiple-testing correction to identify significant differences in mitochondrial perimeter size. A one-way ANOVA was used with Holm-Sidak multiple-testing correction for the OPA1 data after normalization to control. A two-way ANOVA with Holm-Sidak multiple-testing correction was used to compare CL levels between control and each patient sample. A *p* value < 0.05 was considered significant for all analyses.

## Results

### Generation of iPSCs and Identification of *DNAJC19* Mutation

Control and DCMA iPSCs were generated from PBMCs using non-integrative Sendai virus reprogramming.^14^ The iPSCs used in this study came from two Hutterite children from different families (Table 1) harboring the identical homozygous *DNAJC19* intronic pathogenic variant NG_022933.1:c.130-1G>C (rs137854888), that appears to be unique to the Hutterite population in southern Alberta.^2^ The control iPSC line was derived from a healthy, non-Hutterite individual. The generated iPSCs were characterized using standard pluripotency tests.^20^ Immunofluorescence analysis of the iPSCs confirmed positive expression of the human embryonic stem cell (hESC) marker proteins SSEA4 and OCT4 (Supplemental Fig. S1).

Through Sanger sequencing of gDNA isolated from the iPSCs, we demonstrated the presence of the G/C transversion at the conserved AG splice acceptor site of intron 3, confirming the presence of the causative *DNAJC19* IVS3-1G>C mutation (Fig. 1A). Further, to confirm the identity of the cDNA product, we sequenced the 445 bp and 525 bp cDNAs from DCMA and control iPSCs, respectively. As expected, the 445 bp cDNA from the patient iPSCs lacked the coding sequence corresponding to exon 4, while the 525 bp cDNA from control-iPSCs contained the full coding sequence of *DNAJC19* (Supplemental Fig. S2).

**Figure 1.**
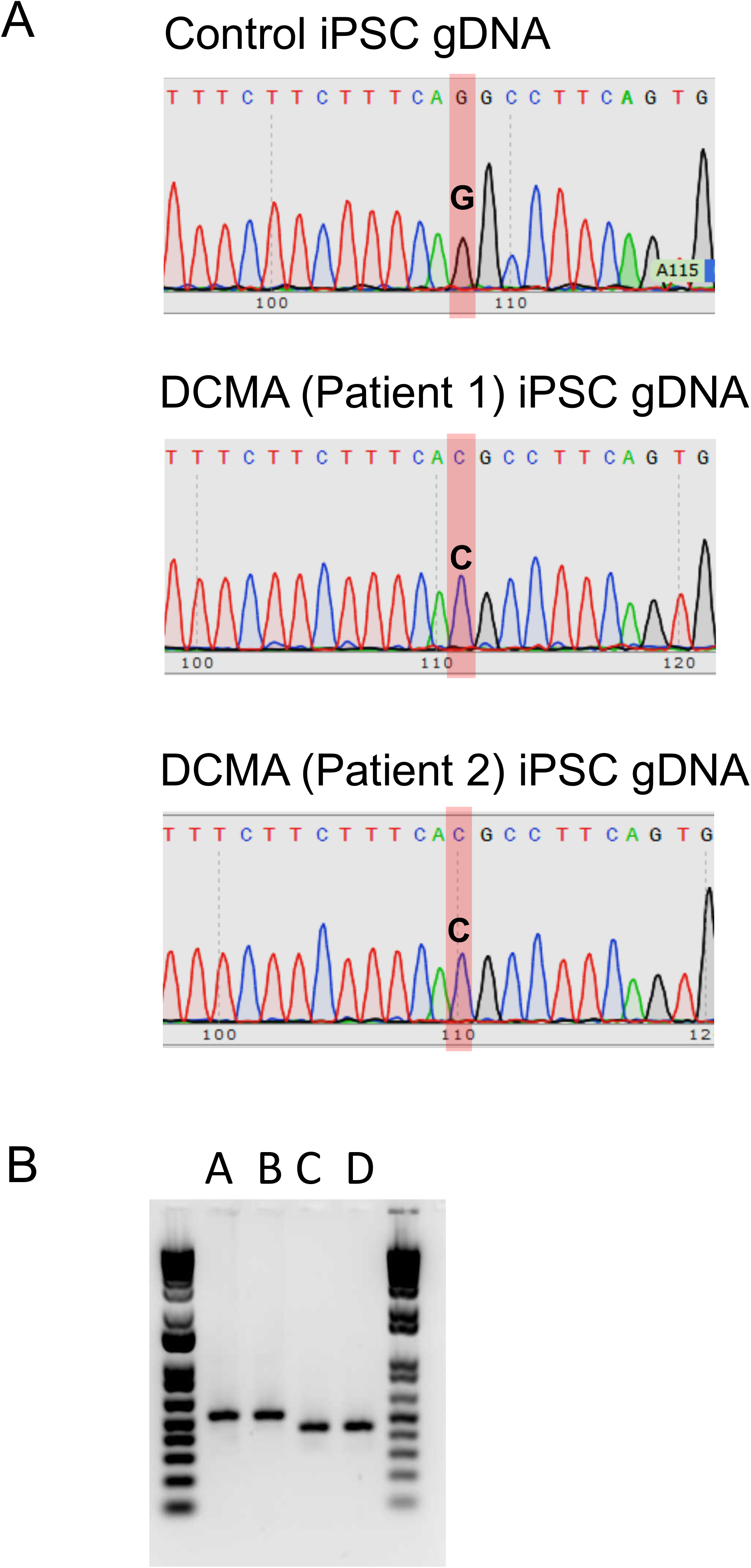
*DNAJC19* mutation and abnormal splicing in DCMA iPSCs. (**A**) Sequencing chromatograms of genomic DNA (gDNA) isolated from control and DCMA iPSCs showing the causative homozygous *DNAJC19* mutation (IVS3-1 G>C) in the DCMA iPSCs. (**B**) RT-PCR analysis of the *DNAJC19* gene shows a truncated product (445 bp) in iPSCs from patient 1 and patient 2 (lanes C and D) compared to the 525 bp product from wild-type hESC and control-iPSCs (lanes A and B) due to the intronic mutation.

We next confirmed the presence of the *DNAJC19* intronic splicing mutation rs137854888 in the patient iPSCs. The iPSCs from both Patient 1 and Patient 2 showed truncated RT-PCR products (445 bp) compared to wild-type hESCs and control-iPSCs (525 bp) (Fig. 1B). These results indicate that we successfully established patient-specific DCMA iPSCs that carry the pathogenic *DNAJC19* mutation.

### Differentiation and Characterization of DCMA iPSC-CMs

To efficiently generate iPSC-CMs for DCMA modeling, we employed a modified monolayer-based myocardial differentiation protocol.^16^ Confluent iPSC colonies from both control and patient groups were treated with the GSK3 inhibitor CHIR99021 followed by inhibition of Wnt signaling by IWP-4 (Supplemental Fig. S3). Corroborating previously published reports, we found that sequential inhibition of the GSK3 and Wnt signaling pathways stimulated cardiogenesis and caused early formation of cardiac tubes and cardiac mesoderm between days 3-5 of differentiation.^16^ Cardiac progenitors and maturation of cardiac tube-like structures were observed on days 5-8 of differentiation. The first cluster of beating cells was observed on days 7–8 after the induction of differentiation. Following lactate-based metabolic selection,^17^ robustly contracting cardiomyocytes were visible on day 12 post-differentiation and by day 20 or 21, cardiomyocytes were obtained for further analysis (Supplemental Movie 1). Characterization of the iPSC-CMs from day 20 showed cytoplasmic expression of TNNT2 and nuclear expression of the early mesodermal marker NKX2.5 (Supplemental Fig. 3B). We next confirmed the efficiency of our cardiac differentiation protocol through flow cytometry analysis selecting for TNNT2-positive cells. For the control iPSC-CMs, 92.8% were TNNT2-positive and the DCMA iPSC-CMs were 90.1% and 69.2% TNNT2-positive for Patients 1 and 2, respectively (Supplemental Figs. 3C and 3D). Our differentiation protocol enabled efficient production of pure cardiomyocytes but the observation of some cells with round, polygonal and rod-shaped morphologies suggest the residual presence of some immature phenotypes. Further approaches to optimize iPSC-CM maturation could be useful to overcome this limitation of our model system.^21^

### Cardiolipin Composition

Multiple CL species in iPSCs and iPSC-CMs were quantified using mass spectrometry. Several small but statistically significant differences were seen between the control and patient iPSCs (Fig. 2 and Supplemental Table S2). Fewer CL species were detectable in the iPSCs compared to the iPSC-CMs (14 vs. 21) and, importantly, the mature form, CL(18:2)4, was not present in significant amounts (<1%) in the iPSCs. In the iPSCs, CL(18:2)(18:1)3 predominanted making up ≥40% of the total CL. In the iPSC-CMs, no consistent and significant differences between the DCMA patients and the control strain were seen (Fig. 3 and Supplemental Table S3). There was greater CL diversity in the iPSC-CMs and mature CL contributed between 6.5-14.4% of the total.

**Figure 2.**
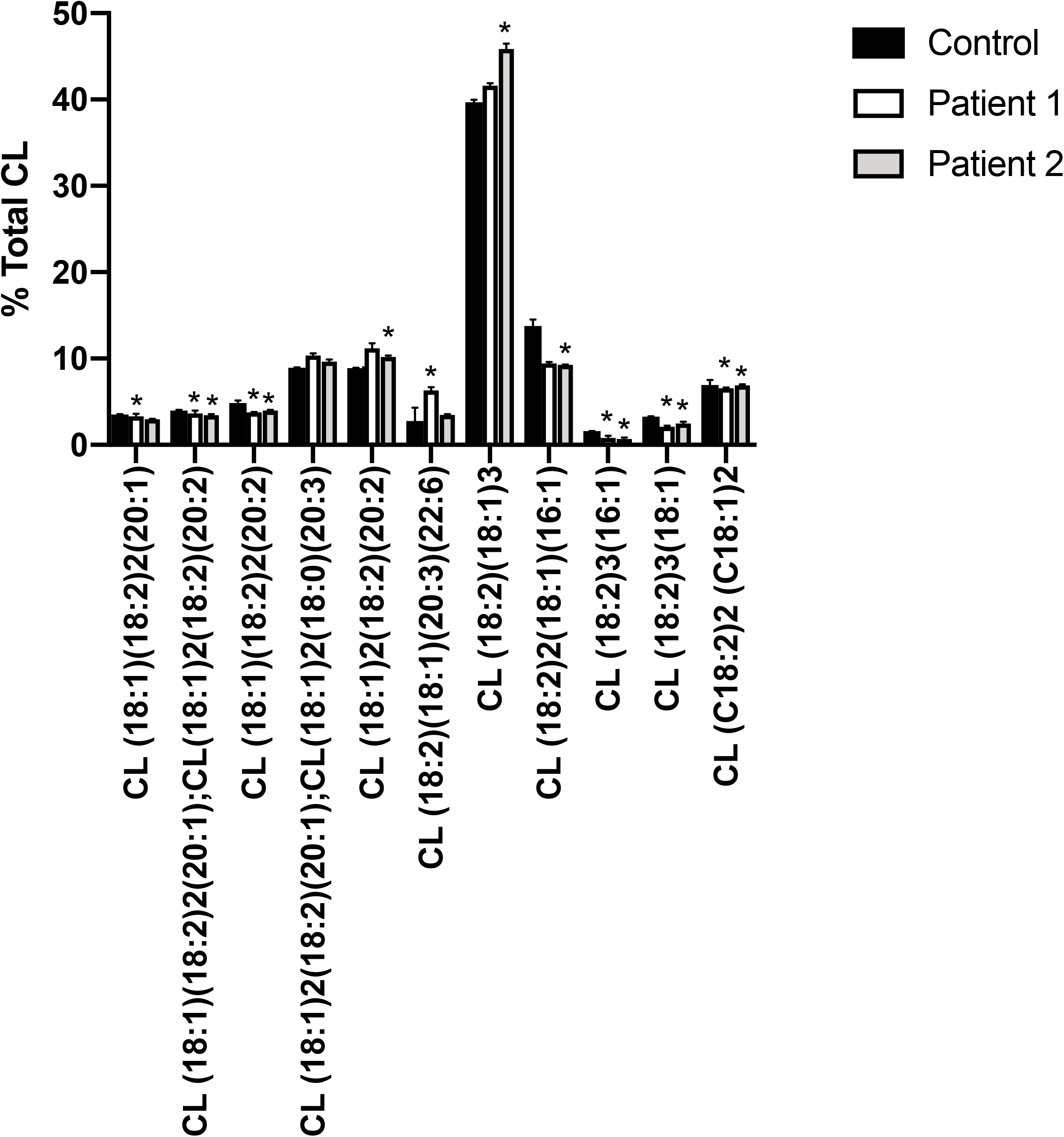
Cardiolipin composition (as a percentage of total cardiolipin) for control and DCMA iPSCs. Of the potential 36 molecular species detectable, only those >1% are shown. *Indicates those patient samples that were significantly different (p<0.05) from control as determined by a two-way ANOVA. Data represent the mean ± SD of two separate cell preparations. The complete dataset is presented in Supplemental Table S2.

**Figure 3.**
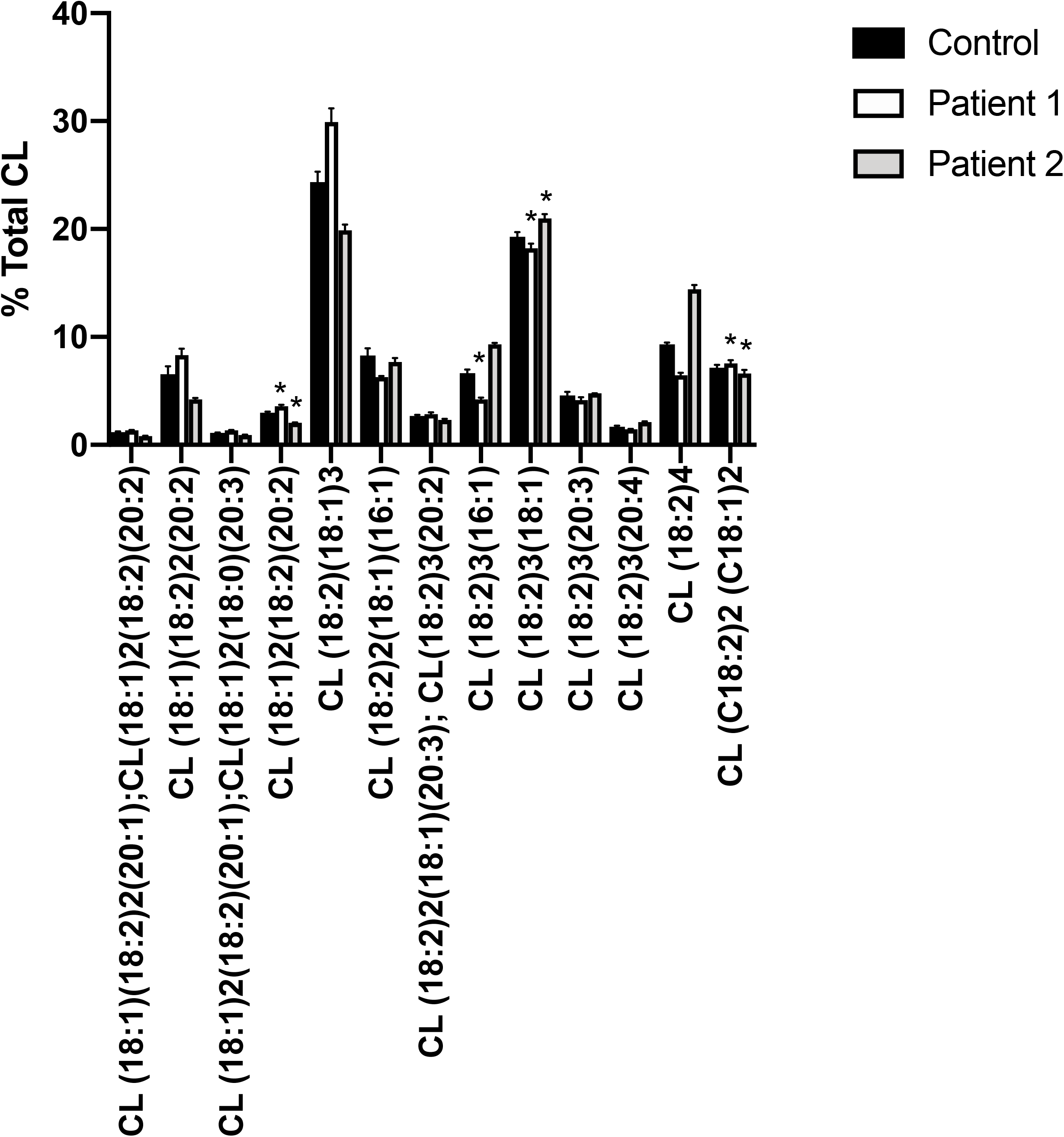
Cardiolipin composition (as a percentage of total cardiolipin) for control and DCMA iPSC-CMs cultured for 21 days. Of the potential 36 molecular species detectable, only those >1% are shown. *Indicates those patient samples that were significantly different (p<0.05) from control as determined by a two-way ANOVA. Data represent the mean ± SD of three separate cell preparations. The complete dataset is presented in Supplemental Table S3.

### Mitochondrial Fragmentation in DCMA iPSC-CMs is Reversible by SS-31

To determine whether the *DNAJC19* mutation affected mitochondrial structure in DCMA iPSC-CMs, we stained both control and patient-derived iPSC-CMs for the outer mitochondrial membrane (OMM) marker TOMM20. Qualitatively, mitochondrial networks in DCMA were highly fragmented compared to the control (Fig. 4A). The mitochondrial networks in DCMA appeared less connected and mitochondria were shorter, whereas control cells showed more fused mitochondrial networks. We then tested whether the mitochondrially-targeted peptide SS-31 could restore mitochondrial morphology and, indeed, the networks in DCMA iPSC-CMs appeared to be less fragmented following SS-31 treatment (Fig. 4A).

**Figure 4.**
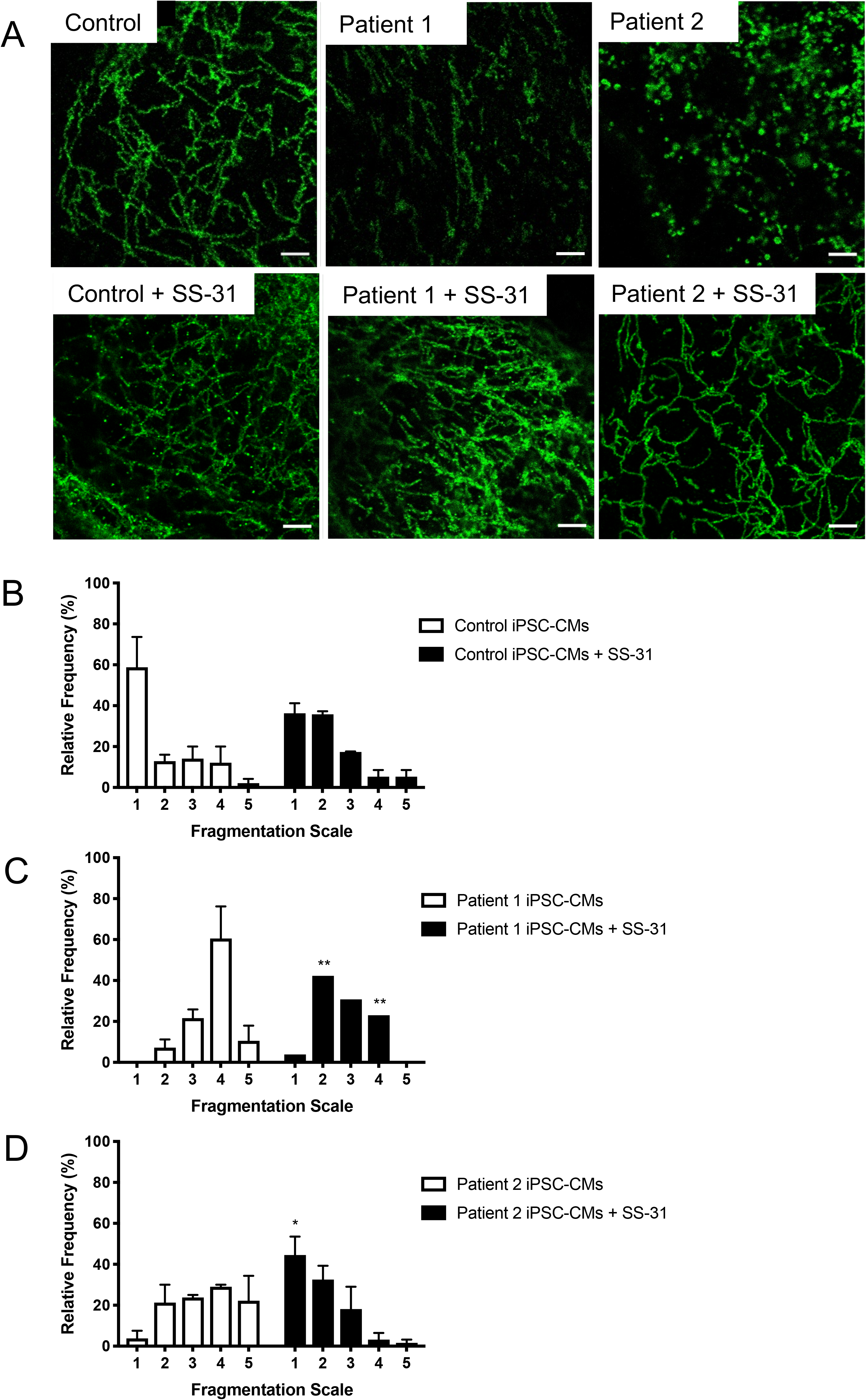
Mitochondrial morphology of iPSC-CMs before and after SS-31 treatment as visualized by TOMM20 immunostaining. (**A**) Representative confocal microscopy images of mitochondria from untreated iPSC-CMs and after incubation with SS-31 for 24 hours. Scale bar 20 μm. (**B-D**) Manual quantification of mitochondrial fragmentation in iPSC-CMs with and without SS-31. Mitochondrial morphology was classified into 5 groups: hyper-fused (grade 1), intermediate (grades 2, 3, 4) and fully fragmented (grade 5). Data are presented as mean ± SEM from two separate blinded manual evaluations of 30 cells quantified per cell line and per condition and compared to untreated using ANOVA. (**B**) Histograms depicting relative frequency of fragmentation categories in control iPSC-CMs with and without SS-31. (**C**) Histograms depicting categories of fragmentation in DCMA iPSC-CMs (Patient 1) with and without incubation with SS-31. **p< 0.01. **(D)** Histograms depicting relative frequency of fragmentation in DCMA iPSC-CMs (Patient 2) with and without with SS-31. *p = 0.01.

Manual quantification, based on a previously reported grading scale,^18^ confirmed our observation of significantly greater fragmentation of the mitochondrial network in both DCMA patients compared to control with a shift towards more fragmented mitochondrial networks in the patient strains (Fig. 4B-D). However, in the presence of SS-31, the DCMA iPSC-CMs demonstrated a significant shift towards a less fragmented phenotype (Figs. 4C and 4D). In contrast, SS-31 treatment did not have any significant effect on control iPSC-CMs (Fig. 4A). The relative frequencies of the fragmentation scale assignments before and after SS-31 are listed in Supplemental Table S4. These results demonstrate the presence of highly fragmented mitochondria in DCMA-derived iPSC-CMs associated with the *DNAJC19* mutation and the reversal of this phenotype by the peptide SS-31.

### Reversal of Defects in Mitochondrial Ultrastructure in DCMA iPSC-CMs by SS-31

We next used TEM to evaluate the effects of the *DNAJC19* mutation on the mitochondrial ultrastructure of our DCMA iPSC-CMs. Consistent with our observations of mitochondrial network fragmentation, we found abnormally-shaped, swollen mitochondria with disorganized cristae in DCMA compared to elongated mitochondria with well-defined cristae in control (Fig. 5A). Perimeter length of mitochondria, a measure of mitochondrial size, in the DCMA iPSC-CMs was significantly smaller (p<0.01 for Patient 1 and p<0.05 for Patient 2) compared to control cells (Fig. 5B). Corroborating the results of our immunocytochemistry, TEM revealed improvements in cristae morphology, mitochondrial shape (Figure 5A) and perimeter length (Fig. 5C) in DCMA iPSC-CMs after SS-31 treatment. These findings indicate that the *DNAJC19* mutation is also associated with significant changes in the mitochondrial ultrastructure and that these changes can also be reversed by SS-31.

**Figure 5.**
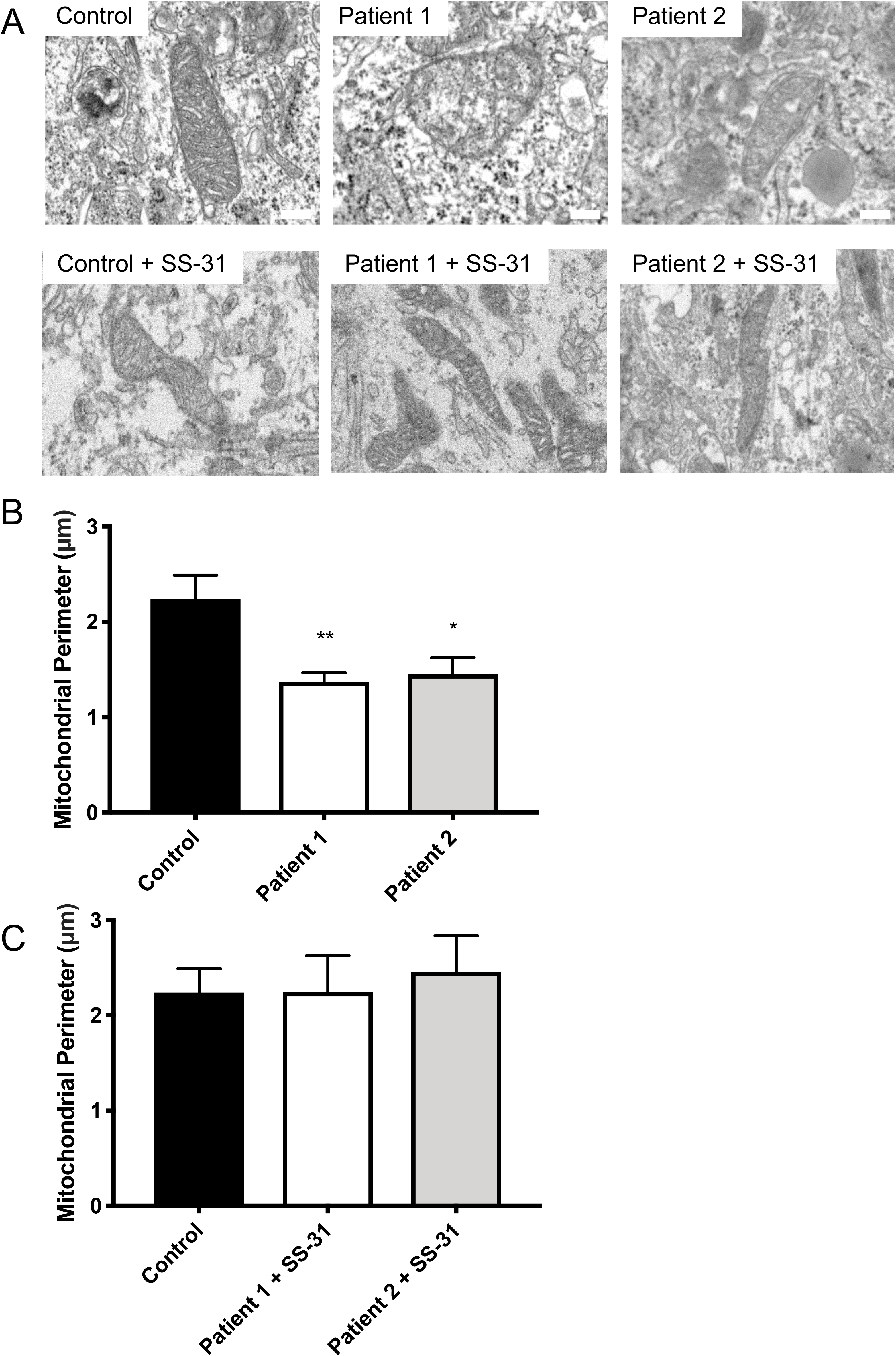
Fragmented mitochondrial ultrastructure in DCMA is normalized by treatment with SS-31. (**A**) Representative TEM images of mitochondria from control and DCMA iPSC-CMs before and after SS-31 treatment. Scale bar 500 nm. (**B**) Quantification of mitochondrial perimeter length in control and DCMA iPSC-CMs without SS-31 treatment. The length of 10 mitochondria was measured per strain and condition. Data are mean ± SEM for two separate experiments. Groups were compared using ANOVA. * p < 0.05, ** p < 0.01. (**C**) Quantification of mitochondrial perimeter length in control and DCMA iPSC-CMs after incubation with SS-31. There were no significant differences between the groups by ANOVA.

### Modulation of Imbalanced OPA1 by SS-31

The appropriate balance between mitochondrial fusion and fission is one of the hallmarks of mitochondrial health.^22–24^ To assess whether the *DNAJC19* mutation affected this balance, we measured the ratio of the long pro-fusion (L-OPA1) and short pro-fission (S-OPA1) isoforms of OPA1, an IMM protein critical for the regulation of mitochondrial fission and fusion. Consistent with the changes in mitochondrial morphology, we found lower levels of L-OPA1 in DCMA iPSC-CMs compared to the control (Fig. 6A), creating a significantly lower ratio of L-OPA1 / S-OPA1 (p<0.01) (Fig. 6C). Treatment with SS-31 reduced the processing of OPA1 to the shorter form in DCMA-iPSC-CMs (Fig. 6B), restoring the isoform ratio back to control levels (Fig. 6D). These results suggest that the *DNAJC19* mutation results in imbalanced OPA1 processing, which interferes with the balance between mitochondrial fusion and fission, resulting in fragmented mitochondria. Restoration of the appropriate OPA1 isoform balance by SS-31 highlights a potential role for this peptide in affecting the abnormal mitochondrial structure seen in DCMA.

**Figure 6.**
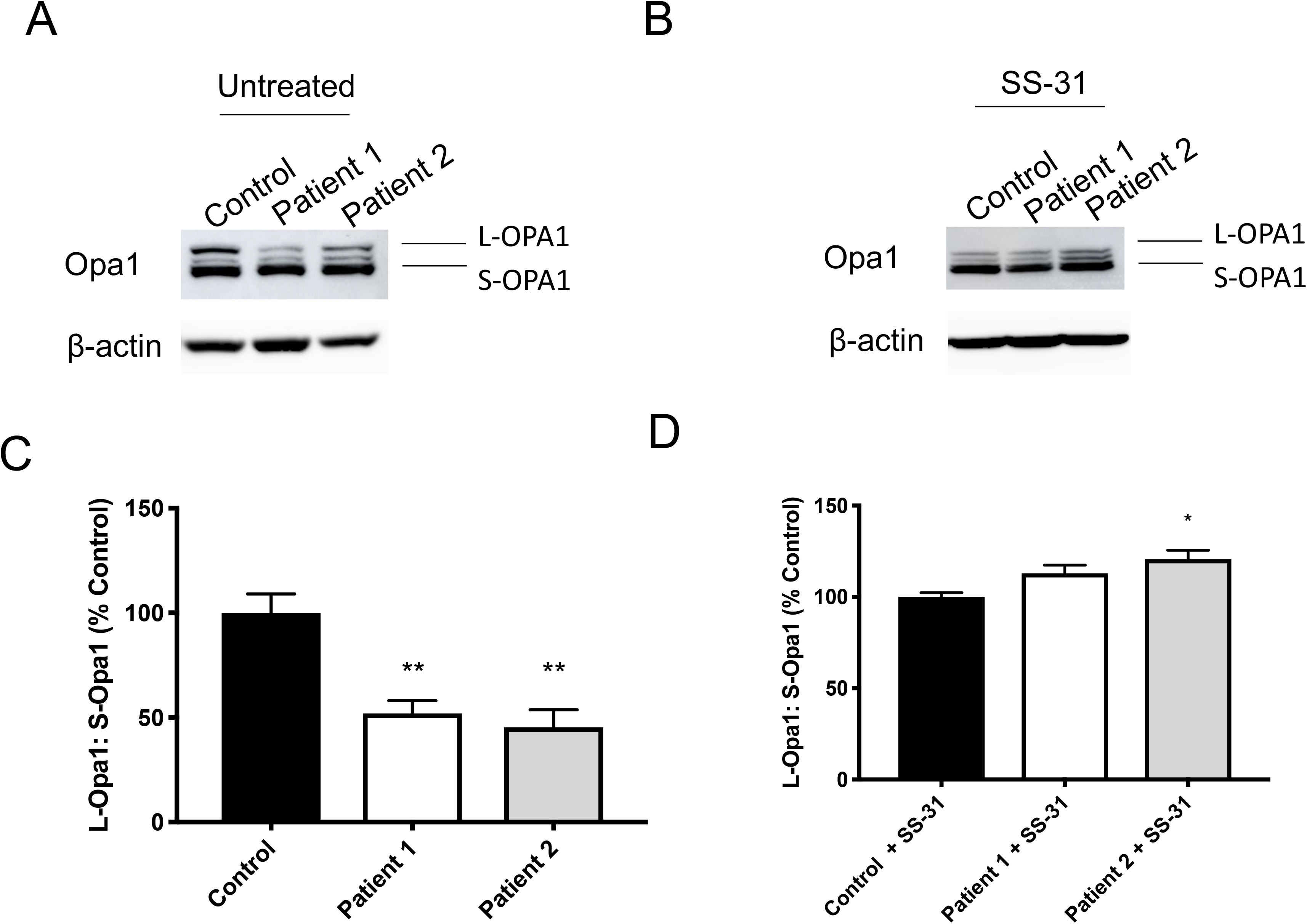
Isoforms of OPA1 protein in iPSC-CMs before and after SS-31 treatment. (**A**) Western blot of control and DCMA iPSC-CMs for OPA1 protein. The long (L-OPA1) and short (S-OPA1) isoforms of OPA1 are shown with β-actin as a loading control. (**B**) Western blot of control and DCMA iPSC-CMs for OPA1 protein after incubation with SS-31 (**C**) Densitometric quantification of Western blots for OPA1 isoforms in control and DCMA iPSC-CMs. Data are shown as percentage of control (n = 4) compared using ANOVA. **p < 0.01. (**D**) Densitometric quantification of Western blots for the OPA1 isoforms in control and DCMA iPSC-CMs after incubation with SS-31. Data are shown as percentage of control (n = 3) compared using ANOVA. *p < 0.05.

## Discussion

In the present study, we describe a novel and unique *in vitro* model for the mitochondrial disease DCMA created by generating patient-specific iPSCs from two children with genetically-and biochemically-confirmed DCMA and severe cardiac dysfunction, followed by the successful differentiation of these cells into beating iPSC-CMs. We confirmed the presence of the causative *DNAJC19* splicing mutation (rs137854888, IVS3-1G>C) in the patient-derived iPSCs which resulted in the loss of the full-length *DNAJC19* transcript. This mutation caused significant mitochondrial fragmentation that was reversible by SS-31.

Although DCMA is purported to be related to Barth syndrome due to similar abnormalities in metabolism and development, we did not observe consistent, significant changes in CL composition, as has been reported for Barth syndrome.^25, 26^ This preliminary observation is of great interest and suggests that the fundamental mechanisms underlying these closely related diseases may be different. However, given the similar phenotypes for DCMA and Barth syndrome, there may exist a cellular relationship between TAZ and DNAJC19 that remains to be elucidated.

DNAJC19 is localized to the IMM but its precise cellular function has not been confirmed in patient-derived cells. Through both confocal microscopy and TEM analyses, we identified highly fragmented mitochondrial networks and ultrastructural defects in DCMA cells confirming that mutated *DNAJC19* leads to impaired mitochondrial structure. In a related observation, we found an imbalance in the isoforms of OPA1 in DCMA iPSC-CMs. The reduction of L-OPA1 relative to S-OPA1 in DCMA supports our hypothesis that the *DNAJC19* mutation affects this balance and causes fragmented mitochondria. It has been suggested that DNAJC19 interacts with prohibitin 2 (PHB2).^3^ The loss of the PHB complex, as a consequence of decreased DNAJC19, could lead to the activation of IMM proteases and increased processing of L-OPA1.^27^ It has previously been shown that imbalanced OPA1 processing and mitochondrial fragmentation are associated with heart failure which is a common clinical phenotype in DCMA.^28^

Prior reports have shown that SS-31 interacts specifically with CL to prevent peroxidative damage and maintain mitochondrial structure and integrity.^11–13^ Paralleling results seen in other mitochondrial disorders,^29, 30^ the peptide SS-31 showed promise for its ability to reverse the mitochondrial abnormalities found in DCMA. We found reversible mitochondrial fragmentation, improved mitochondrial ultrastructure and restoration of the OPA1 ratio in DCMA iPSC-CMs following exposure to SS-31. Together, these suggest a potential role for SS-31 as a novel therapeutic for the treatment of DCMA. Although SS-31 improved mitochondrial structure and the balance of OPA1 isoforms in our DCMA-derived iPSC-CMs, the precise mechanism of its action has not been elucidated in this observational study.

## Conclusions

In summary, using a novel *in vitro* cardiomyocyte model we have described the abnormal mitochondrial structure and ultrastructure in DCMA and identified an imbalance of OPA1 isoforms as a possible mechanism or consequence. We have also demonstrated reversal of this phenotype through treatment with SS-31. Our findings suggest that further evaluation of SS-31 in DCMA is warranted and studies into the mechanism of this poorly understood and heterogeneous disease are now feasible using our patient-derived cells.

## Funding Sources

This work was supported by research grants from the Children’s Cardiomyopathy Foundation and the Alberta Children’s Hospital Foundation to S.C.G.

## Author Contributions

L.R., P.M., A.K., T.S., D.R. and S.C.G. conceived and designed experiments. J.C.W. reprogrammed the PBMCs into iPSCs and validated pluripotency. G.M. and L.R. optimized the iPSC-CM differentiation protocol. L.R., P.M., R.S., S.L., T.Z. performed all experiments. A.R. performed the CL analysis. L.R., P.M., R.S. and F.I. performed the data analysis. L.R., P.M., D.R. and S.C.G wrote the manuscript. All authors read and approved the manuscript.

## Acknowledgements

We would like to acknowledge the imaging resources of the Charbonneau Microscopy Facility and the Microscopy and Imaging Facility at the University of Calgary. We would also like to thank the Flow Cytometry Core Facility at the University of Calgary for data processing.

## Data availability

The datasets generated and analyzed for this study are available from the corresponding author on reasonable request.

